# PIF4/HEMERA-mediated daytime thermosensory growth requires the Mediator subunit MED14

**DOI:** 10.1101/2022.03.02.482654

**Authors:** Abhishesh Bajracharya, Jing Xi, Karlie F. Grace, Eden E. Bayer, Chloe A. Grant, Caroline H. Clutton, Scott R. Baerson, Ameeta K. Agarwal, Yongjian Qiu

## Abstract

While moderately elevated ambient temperatures do not trigger stress responses in plants, they do significantly stimulate the growth of specific organs through a process known as thermomorphogenesis. The basic helix-loop-helix transcription factor PHYTOCHROME-INTERACTING FACTOR 4 (PIF4) plays a central role in regulating thermomorphogenetic hypocotyl elongation in various plant species, including *Arabidopsis thaliana*. Although it is well known that PIF4 promotes plant thermosensory growth by activating genes involved in the biosynthesis and signaling of the phytohormone auxin, the detailed molecular mechanism of such transcriptional activation is not clear. Our previous studies demonstrated that HEMERA (HMR), a transcriptional co-activator of PIF4, promotes the thermo-induced expression of PIF4 target genes through its nine-amino-acid transactivation domain (9aaTAD). In this report, we investigate the role of the Mediator complex in the PIF4/HMR-mediated thermoresponsive gene expression. Through the characterization of various mutants of the Mediator complex, a tail subunit named MED14 is identified as an essential factor for thermomorphogenetic hypocotyl growth. MED14 is required for the thermal induction of PIF4 target genes but has a marginal effect on the levels of PIF4 and HMR. Further transcriptomic analyses confirm that the expression of numerous PIF4/HMR-dependent, auxin-related genes requires MED14 at warm temperatures. Moreover, PIF4 and HMR physically interact with MED14 and both are indispensable for the association of MED14 with the promoters of these thermoresponsive genes. Taken together, these results unveil an important thermomorphogenetic mechanism, in which PIF4 and HMR recruit the Mediator complex to activate auxin-related growth-promoting genes when plants sense moderate increases in ambient temperature.

**One-sentence summary:** The Mediator subunit MED14 promotes thermomorphogenesis by interacting with PHYTOCHROME-INTERACTING FACTOR 4 and HEMERA to induce the expression of growth-promoting genes at elevated temperatures.

## Introduction

Plants are highly sensitive to environmental temperature changes. Moderately elevated ambient temperatures below heat shock (between 12 °C and 27 °C) do not trigger stress responses, but they may drastically alter plant growth and development, including rapid stem and root elongation, enhanced petiole hyponastic growth, early flowering, and reduced stomatal index. Collectively, these responses are referred to as thermomorphogenesis (Casal and Balasubramanian, 2019; Ludwig et al., 2021; Park et al., 2021).

Thermomorphogenetic responses, such as thermosensory hypocotyl elongation, involve massive transcriptomic reprogramming, as demonstrated in the model dicotyledonous plant *Arabidopsis thaliana* (Liu and Charng, 2013; Sidaway-Lee et al., 2014; Cortijo et al., 2017; Ding et al., 2018; Han et al., 2019; Bordiya et al., 2020; Jin et al., 2020; Lee et al., 2020; Lee et al., 2021b; Lee et al., 2021c). A major effect of this transcriptomic reprogramming is to activate genes involved in auxin synthesis, such as *YUCCA8* (*YUC8*), and auxin signaling, such as *INDOLE-3-ACETIC ACID INDUCIBLE 19* & *29* (*IAA19* & *IAA29*) (Franklin et al., 2011; Sun et al., 2012; Ma et al., 2016; Zhu et al., 2016). PHYTOCHROME-INTERACTING FACTOR 4 (PIF4), a central transcriptional regulator of thermomorphogenesis (Qiu, 2020), activates these auxin-related genes in response to warm temperatures (Franklin et al., 2011; Sun et al., 2012; Ma et al., 2016; Zhu et al., 2016). PIF4 belongs to an eight-member basic Helix-Loop-Helix (bHLH) transcription factor family, which was first identified as a key player in transducing light signals perceived by the red/far-red photoreceptors known as phytochromes (Leivar and Quail, 2011). PIFs are integrators of light and various environmental and developmental signals (Leivar and Quail, 2011; Leivar and Monte, 2014; Pham et al., 2018). Although PIFs show apparent functional redundancy in multiple morphogenetic responses (Leivar and Monte, 2014; Pham et al., 2018), thermomorphogenesis is mediated primarily through PIF4 (Koini et al., 2009; Stavang et al., 2009; Kumar et al., 2012; Qiu et al., 2019). As an early ambient temperature signaling component, PIF4 is regulated at multiple levels, including transcription, post-translational modification, DNA-binding affinity, transcriptional activity, and protein stability (Qiu, 2020). Recent reports revealed distinct temperature-signaling mechanisms under different light conditions or photoperiod regimes, indicating more complex regulation of PIF4-mediated thermal responses by light quality, day length, and the circadian clock (Box et al., 2015; Park et al., 2017; Qiu et al., 2019).

Besides its role in thermomorphogenesis, PIF4 also activates growth-promoting genes during skotomorphogenesis (growth in darkness) and shade-avoidance responses (Leivar and Monte, 2014). PIF4 possesses a transactivation domain (TAD) that resembles that of PIF3 (Dalton et al., 2016; Yoo et al., 2021). The tomato PIF4 activates shade-induced genes by interacting with MED25, a tail component of the Mediator complex (Sun et al., 2020). These data suggest that PIF4 has a functional TAD that promotes shade-avoidance responses. However, whether PIF4’s TAD is equally important for its activity in thermomorphogenesis is unknown. We previously demonstrated that PIF4-mediated thermomorphogenetic control of growth-promoting gene expression requires the transcriptional coactivator HEMERA (HMR) (Qiu et al., 2019). HMR physically interacts with PIF4 and activates the thermo-inducible PIF4 target genes through an acidic nine-amino-acid transactivation domain (9aaTAD). A weak allele named *hmr-22* harbors a D516N mutation in the 9aaTAD, largely disrupting HMR’s transactivation activity and leading to defects in the thermosensory growth (Qiu et al., 2015; Qiu et al., 2019). Mechanistic insights into the function of HMR’s TAD in regulating PIF4 activity have yet to be revealed.

HMR’s 9aaTAD resembles the TADs present in prototypic acidic transcription activators, such as VP16, GAL4, GCN4, and MYC (Piskacek et al., 2007). These acidic TADs interact directly with subunits of the Mediator complex to facilitate the assembly or the stability of the RNA polymerase II (Pol II) preinitiation complex (PIC) at the transcriptional initiation site (Borggrefe and Yue, 2011; Vojnic et al., 2011). The Mediator complex is a multi-subunit transcriptional coordinator that transmits signals from transcription activators to the Pol II preinitiation transcription activators and is required for the transcription of virtually all Pol II-transcribed genes (Dolan et al., 2017).

Much of our current understanding of the three-dimensional structure of the Mediator complex has come from studies performed in yeast and mammals (Robinson et al., 2016; Nozawa et al., 2017; Schilbach et al., 2017; Tsai et al., 2017; Chen et al., 2021; Rengachari et al., 2021). Despite the low sequence similarity between some Arabidopsis MED subunits and those of yeast and metazoans, more than 20 MED subunits are conserved across yeast, plants, and metazoans (Bourbon, 2008). Furthermore, the Mediator complexes in all three Kingdoms share a generally similar modular composition, with subunits organized into head, middle, tail, and kinase subcomplexes (Malik and Roeder, 2010; Buendía-Monreal and Gillmor, 2016; Dolan and Chapple, 2017). The head and middle modules form the essential core Mediator that directly interacts with PIC (Nozawa et al., 2017; Schilbach et al., 2017). PIC-Mediator assembly further creates a Head-Middle sandwich to stabilize two Pol II C-terminal domain (CTD) segments and position CTD for phosphorylation (Chen et al., 2021). In contrast to the head and middle subunits, the subunits of the tail module are relatively loosely associated with each other and are targeted by activators and repressors (Malik and Roeder, 2010; Cevher et al., 2014; Zhao et al., 2021).

To further elucidate the molecular mechanism by which PIF4 and HMR recruit the transcriptional machinery when activating the expression of thermoresponsive genes, we performed a reverse genetic screen for Arabidopsis Mediator mutants that show reduced sensitivity to moderate temperature elevations. Here, we report the identification of MED14 as a key subunit of the Mediator complex that is recruited by PIF4 and HMR to activate auxin-related, growth-promoting genes in response to warm temperatures.

## Results

### MED14 is required for daytime thermosensory hypocotyl growth

To investigate the role of the Mediator complex in thermomorphogenesis and search for Mediator subunits that work with PIF4 and HMR to induce the transcription of thermo-responsive genes, we first tested the thermal sensitivity of mutants of various Mediator subunits. Most mutants of the Mediator subunits in the head, middle, and kinase modules showed similar responses to the warm temperature as wild type—the ratio of hypocotyl length between 27 °C and 20 °C were comparable with those in the wild-type seedlings (Supplemental Figure S1). In comparison, mutants of several tail subunits, including MED14, MED16, and MED25, were hyposensitive to the increased temperature (Figure 1A-C). The ratios of hypocotyl length at 27 °C to that at 20 °C in *med14* (SAIL_373_C07), *med16-2* (SALK_048091), *med16-3* (WISCDSLOX504A08), *med25-2* (SALK_129555C), and *med25-3* (SALK_059316C) were significantly lower than those of wild-type seedlings (Figure 1C). Among these mutants, *med14* showed the most dramatic reduction in thermomorphogenetic hypocotyl growth—it had only 37% of the wild-type thermal response and ~70% of the wild-type hypocotyl length at 27 °C.

**Figure 1.**
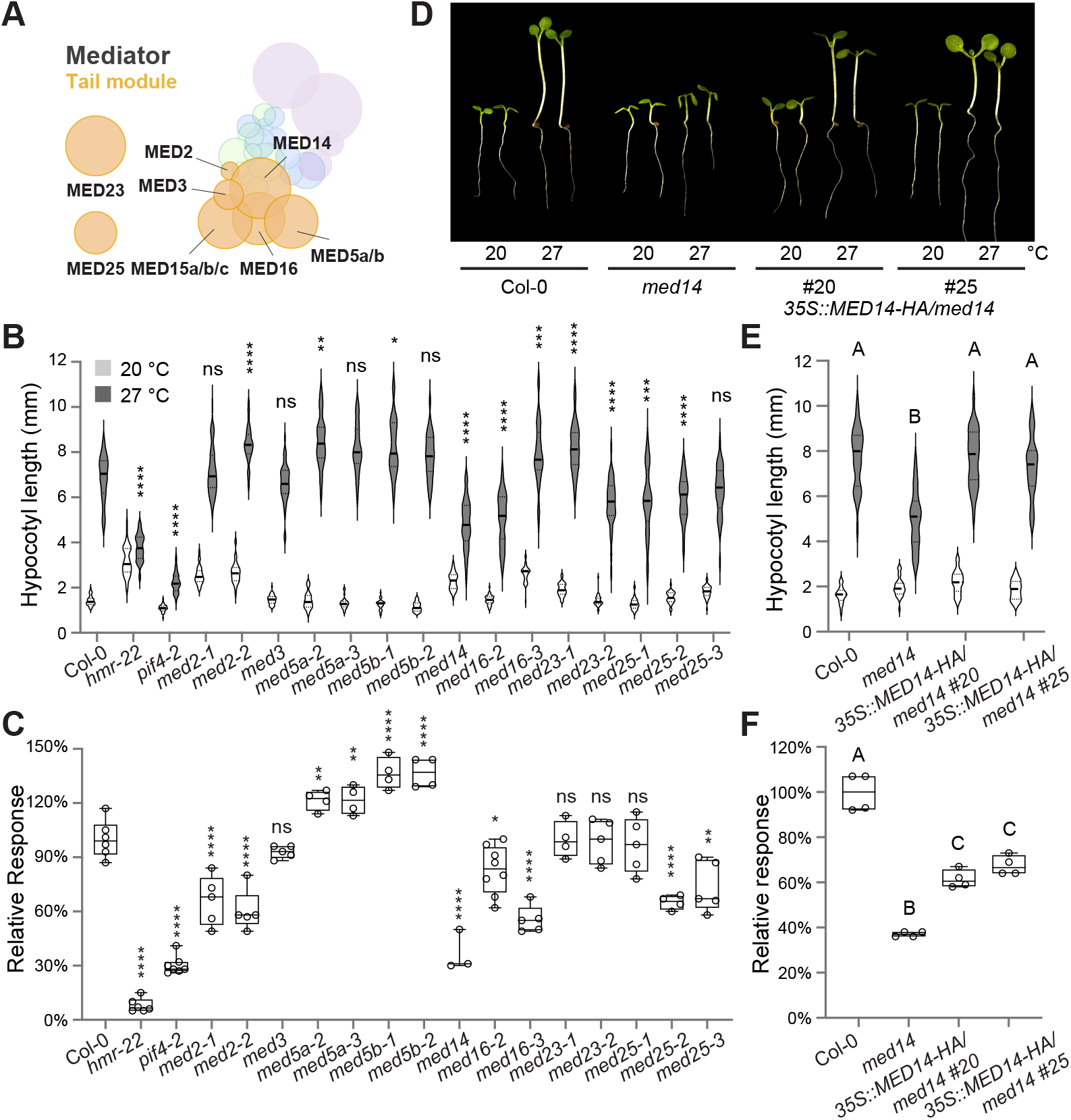
MED14 is a crucial regulator of thermomorphogenesis. (A) Schematic illustration of the Arabidopsis Mediator complex, with an emphasis on components in the tail module. The size of each circle reflects the relative protein size (predicted molecular weight) of each mediator subunit. The relative position of each mediator component is based on the cryo-EM and crystal structures of yeast and human Mediator (Tsai et al., 2014; Robinson et al., 2015) and interaction data of Arabidopsis mediator subunits (Maji et al., 2019). Note that the positions of MED23 and MED25 in the tail module are undetermined. Protein subunits in each module are colored in different colors: head module, green; middle module, blue; tail module, orange; cyclin kinase module, purple. See Supplemental Figure S1 for clear compositions of each module. (B) Hypocotyl length measurements of mutant seedlings of mediator tail subunits. The white and grey bars represent hypocotyl length measurements at 20 °C and 27 °C, respectively. The results of the one-way ANOVA analysis comparing the absolute hypocotyl length between Col-0 and each mutant grown at 27 °C are shown (n > 30). ****, *p* < 0.0001; ***, *p* < 0.001; **, *p* < 0.01; *, *p* < 0.05; ns, not significant (*p* ≥ 0.05). (C) Comparison of the relative thermal response among the seedlings in (B). The relative response is defined as the relative hypocotyl response to 27 °C of a mutant compared with that of Col-0 (which is set at 100%). The results of the one-way ANOVA analysis comparing the relative response between Col-0 and each mutant are shown (n ≥ 3). ****, *p* < 0.0001; ***, *p* < 0.001; **, *p* < 0.01; *, *p* < 0.05; ns, not significant (*p* ≥ 0.05). (D) Representative images of 4-d-old Col-0, *med14*, and *35S::MED14-HA/med14* (#20 and #25) seedlings at either 20 °C or 27 °C. (E) Hypocotyl length measurements of seedlings in (D). Different letters denote significant statistical differences between the absolute hypocotyl length of each line grown at 27 °C (one-way ANOVA, n > 30, *p* < 0.0001). (F) Comparison of the relative thermal response among the seedlings in (D). Different letters denote significant statistical differences between the relative response of each line (one-way ANOVA, n ≥ 4, *p* < 0.0001).

The *med14* mutant (SAIL_373_C07) was the only viable T-DNA insertion line with dramatically reduced *MED14* expression (Zhang et al., 2013). To further confirm that the thermomorphogenetic defect of *med14* is caused by decreased *MED14* levels, we generated *MED14* overexpression lines in the *med14* mutant background. A full-length *MED14* CDS driven by the 35S Cauliflower Mosaic Virus (CaMV) constitutive promoter was expressed with a C-terminal human influenza hemagglutinin (HA) tag. The resulting transgenic plants, designated *35S::MED14-HA/med14*, successfully complemented the morphological phenotype of *med14* at 27 °C and largely rescued its reduced sensitivity in thermomorphogenetic hypocotyl growth (Figure 1D-F). These data suggest that MED14 plays a crucial role in Arabidopsis thermosensory growth.

### MED14 regulates the thermal induction of key auxin-related PIF4 targets

PIF4 and HMR promote thermosensory hypocotyl growth by activating key genes in auxin biosynthesis (e.g., *YUC8*) and signaling (e.g., *IAA19* and *IAA29*). The identification of MED14’s role in thermomorphogenesis prompted us to test whether MED14 is the tail component that works with PIF4 and HMR in activating the expression of these growth-related, thermo-induced genes. Toward this end, we first examined the steady-state transcript levels of *YUC8*, *IAA19*, and *IAA29* in 4-d-old Col-0 and *med14* seedlings grown at lower and warmer temperatures. While the transcription of all three genes was stimulated at 27 °C in Col-0, these increases were not observed in *med14* (Figure 2A). Similarly, a 6-hour warm temperature treatment greatly induced the expression levels of all three markers in Col-0, but not in the *med14* mutant (Figure 2B).

**Figure 2.**
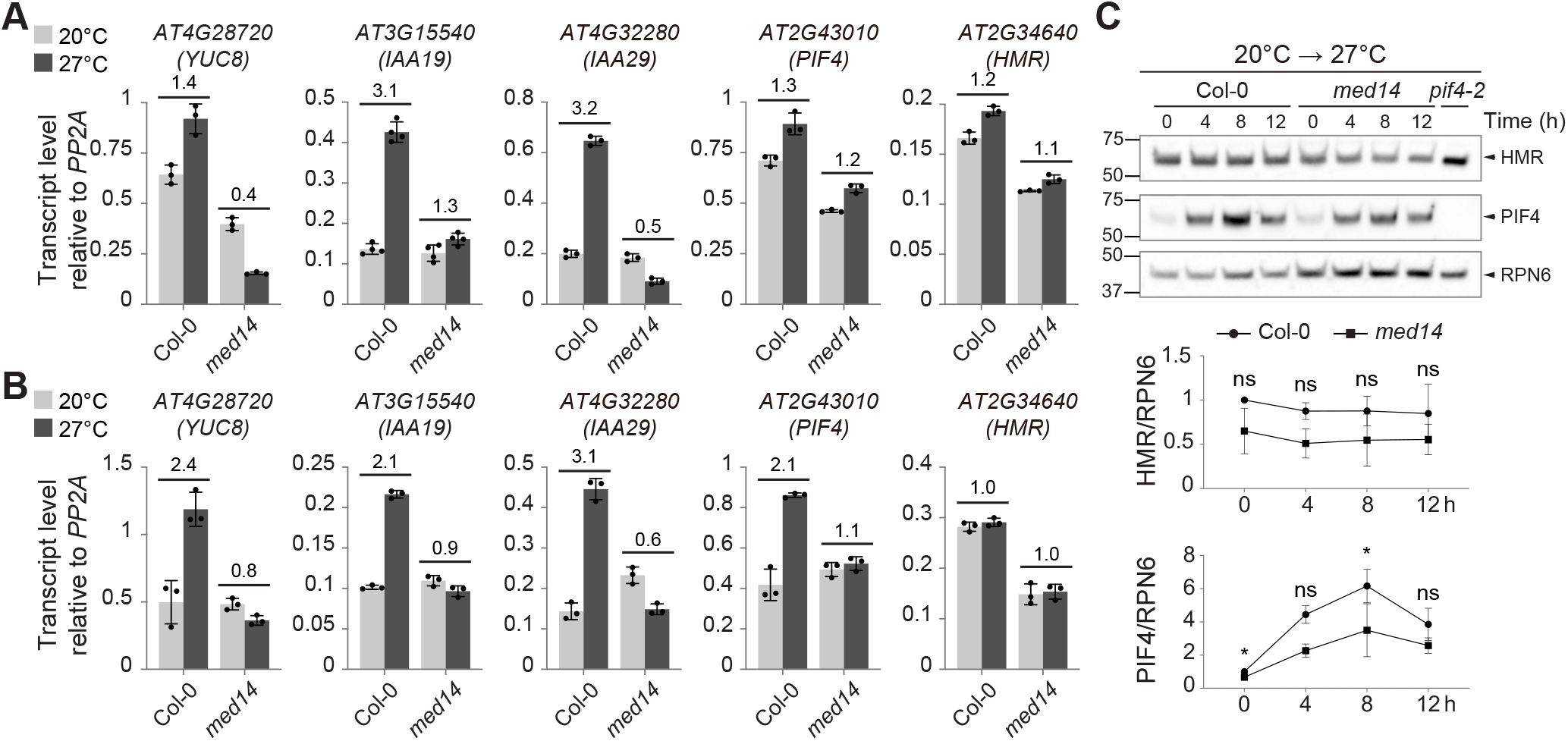
MED14 is required for the thermo-induced expression of PIF4 target genes. (A) RT-qPCR analysis of the steady-state transcript levels of *YUC8*, *IAA19*, *IAA29*, *PIF4*, and *HMR* in Col-0 and *med14* seedlings grown in R light for 96 h at 20 °C and 27 °C. Numbers indicate the fold changes between transcript levels at these two temperatures. (B) RT-qPCR analysis of the steady-state transcript levels of *YUC8*, *IAA19*, *IAA29*, *PIF4*, and *HMR* in Col-0 and *med14* during the 21 °C-to-27 °C transitions. Seedlings were grown in R light for 96 h and then transferred to 27 °C in the same light condition. Samples were taken before (light grey) and 6 h after (dark grey) the 27 °C treatment. Fold changes in the transcript levels by the 27 °C treatment are shown above the columns. For (A) and (B), transcript levels were calculated relative to those of *PP2A*. Error bars represent the SD of three technical replicates. (C) Immunoblot analysis of HMR and PIF4 levels in Col-0 and *med14* during the 21 °C-to-27 °C transitions. Seedlings were grown in R light for 96 h and then transferred to 27 °C in the same light condition. Samples were collected and analyzed at the indicated time points. RPN6 was used as a loading control. The relative levels of HMR and PIF4, normalized to RPN6, are shown underneath the immunoblots. The results of the student t-test analysis are shown (n = 3 or 4). ^*^, *p* < 0.05; ns, not significant (*p* ≥ 0.05).

Given that the Mediator complex is essential for all Pol II-mediated transcription (Jeronimo and Robert, 2017; Soutourina, 2018) and that MED14 is a key component of the Mediator complex (Buendía-Monreal and Gillmor, 2016; Yang et al., 2016), the hypomorphic mutation in *med14* might also affect the expression of *PIF4* and *HMR*. Indeed, the absolute transcript levels of *HMR* and *PIF4* in the *med14* mutant were lower than those in Col-0 at 27 °C (Figure 2A). In addition, the thermo-induced expression of *PIF4*, but not *HMR*, was also compromised in *med14* (Figure 2B). We further examined whether the *med14* mutation also affected the HMR and PIF4 expression at the protein level. While no significant difference in the HMR protein level between Col-0 and *med14* seedlings was observed after the thermal treatment (a 20-to-27 °C transition), the PIF4 protein level was slightly reduced in the *med14* mutant when compared with Col-0 (Figure 2C). These data imply that MED14 might contribute to the thermoresponsive expression of the PIF4/HMR-regulated genes (*YUC8*, *IAA19*, and *IAA29*) in two ways—a direct role by assisting PIF4’s and HMR’s transactivation activity and an indirect role by modulating the levels of *PIF4* and *HMR* when plants are exposed to warmer temperatures.

### MED14, HMR, and PIF4 co-regulate a subset of thermoresponsive genes

Previous transcriptomic analyses by other groups have shown that 4-16% of Arabidopsis genes are differentially regulated by warm temperatures and the expression of a large portion of these thermo-regulated genes requires PIF4 (Ding et al., 2018; Jin et al., 2020; Kim et al., 2020; Lee et al., 2020; Lee et al., 2021b; Lee et al., 2021c). The observation that MED14 is indispensable for the thermal induction of several key PIF4/HMR target genes motivated us to further explore its correlation with PIF4 and HMR in controlling the thermoresponsive gene expression on the genome-wide scale. Therefore, we performed RNA-seq on 4-day-old Col-0, *pif4-2*, *hmr-22*, and *med14* seedlings grown at 20 °C and 27 °C under continuous red light.

When comparing expression levels at 27 °C with those at 20 °C, 828 and 938 genes were induced and repressed at least 2-fold by warmer temperature in Col-0 (FDR *p*-value < 0.001, FPKM > 1), respectively (Figure 3A; Supplemental Dataset 1). These results are in agreement with previous studies in which similar growth conditions were used (Lee et al., 2020; Lee et al., 2021c). For example, 483 of 828 thermo-induced genes (58%) and 550 out of 938 (59%) thermo-repressed genes in the current study overlapped with at least one study reported by Lee et al. 2020 and Lee et al. 2021b (Supplemental Figure S2; Supplemental Dataset 2). The *pif4-2*, *hmr-22*, and *med14* mutants displayed distinct transcriptome profiles compared with Col-0 and a large proportion of genes were uniquely induced or repressed at 27 °C in each mutant (Figure 3A; Supplemental Dataset 1). Even though several genes were also shared between Col-0 and each mutant, 324 and 281 SSTF genes were uniquely induced and repressed in Col-0 at 27 °C, respectively (Figure 3A, Supplemental Figure S3A; Supplemental Dataset 3). We, therefore, named these genes as PIF4/HMR/MED14-dependent thermo-induced and thermo-repressed genes. We performed Gene Ontology (GO) analysis on these two groups of thermo-regulated genes. Notably, many genes involved in growth-related phytohormone signaling (e.g., auxin, brassinosteroid, and ethylene) were upregulated in Col-0 but not in these three mutants, and a number of genes involved in cell wall organization and modification were downregulated only in Col-0 (Figure 3B). This observation is consistent with the thermomorphogenetic hypocotyl growth phenotype of these genotypes (Figure 1B, C).

**Figure 3.**
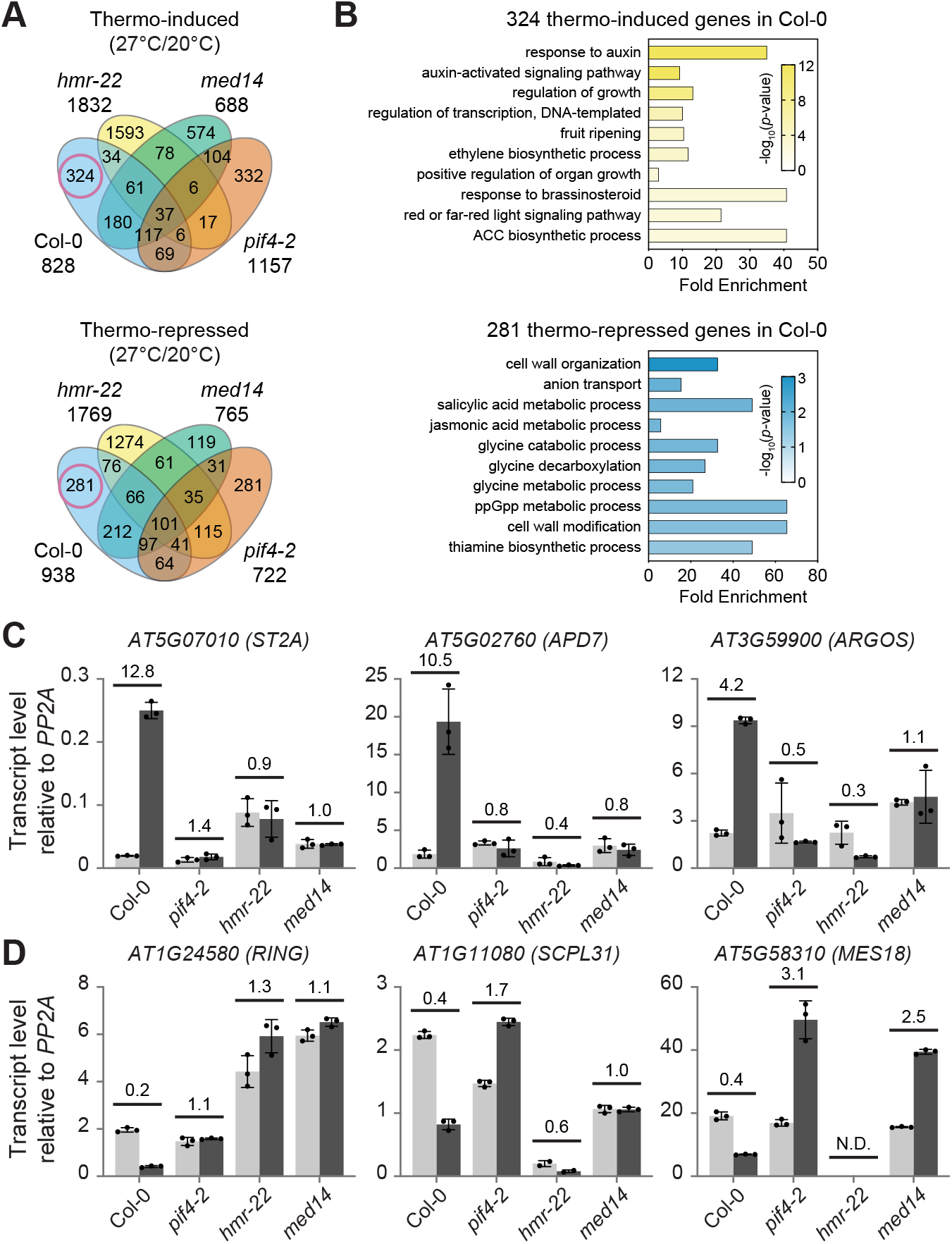
Thermo-induced expression of a group of auxin-related, growth-promoting genes requires PIF4, HMR, and MED14. (A) A Venn diagram shows unique and co-regulated thermo-induced and thermo-repressed genes in wild type (Col-0), *pif4-2*, *hmr-22*, and *med14* mutants. The red circles indicate PIF4/HMR/MED14-dependent genes that were uniquely induced and repressed in Col-0 but not in the mutants at 27 °C. Seedlings were grown in continuous red light for 96 h at either 20 °C or 27 °C, and total RNA was extracted from four biological replicates for RNA-seq analyses. (B) Gene ontology analysis of the 324 PIF4/HMR/MED14-dependent thermo-induced genes and 281 PIF4/HMR/MED14-dependent thermo-repressed genes. The bar represents the fold enrichment and the color indicates the −log10(p-value). ACC, 1-aminocyclopropane-1-carboxylate (an ethylene precursor). ppGpp, guanosine tetraphosphate. (C,D) RT-qPCR analysis of representative PIF4/HMR/MED14-dependent genes in Col-0, *pif4-2*, *hmr-22*, and *med14*. Total RNA was extracted from seedlings grown in continuous red light for 96 h at either 20 °C or 27 °C. The relative expression of three thermo-induced (C) and three thermo-repressed genes (D) were normalized to the expression level of *PP2A* after RT-qPCR. Numbers indicate the fold changes between transcript levels at these two temperatures. N.D., not detected.

To evaluate whether these PIF4/HMR/MED14-dependent thermoresponsive genes are directly or indirectly regulated by PIF4, we compared our datasets with published PIF4-associated genes from three studies (Supplemental Dataset 4). Possibly due to different growth conditions and treatment methods, the number of PIF4-binding genes identified through ChIP-seq assays in the three studies varied from ~1,000 to over 7,000 (Oh et al., 2012; Pfeiffer et al., 2014; Pedmale et al., 2016). We defined the thermo-regulated PIF4 direct targets as the PIF4/HMR/MED14-dependent thermoresponsive genes that were identified in at least two of the three published studies. While less than 10% of the thermo-repressed genes (25 out of 281) were associated with PIF4, about one-third of the thermo-induced genes (114 out of 324) are PIF4 direct targets (Supplemental Figure S3B; Supplemental Dataset 4). Therefore, MED14 may mainly function as a coactivator, rather than a repressor, in regulating thermoresponsive PIF4 targets. GO analyses of these two datasets showed similar results as those of PIF4/HMR/MED14-dependent thermoresponsive genes (Supplemental Figure S3C). Many auxin-responsive genes, including *IAA19*, *IAA29*, and a group of *SMALL AUXIN UP RNA* (*SAUR*) genes, were enriched in the PIF4/HMR/MED14-dependent thermo-induced PIF4 targets; several genes involved in cell wall modification and lignin metabolic process were enriched in PIF4/HMR/MED14-dependent thermo-repressed PIF4 targets (Supplemental Figure S3C).

We further performed RT-qPCR analyses to confirm the RNA-seq results (Figure 3C, D). The expression of three PIF4/HMR/MED14-dependent thermo-induced genes, *ST2A* (AT5G07010), *APD7* (AT5G02760), and *ARGOS* (AT3G59900), were highly induced in Col-0 at 27 °C but showed similar levels at lower and warmer temperatures in *pif4-2*, *hmr-22*, and *med14* mutants (Figure 3C). The expression of three PIF4/HMR/MED14-dependent thermo-repressed genes, *RING* (AT1G24580), *SCPL31* (AT1G11080), and *MES18* (AT5G58310), were greatly reduced in Col-0 at 27 °C but not in those three mutants (Figure 3D). Taken together, these data suggest that MED14 co-regulates a group of growth-related, thermoresponsive genes with PIF4 and HMR.

### PIF4 and HMR physically interact with MED14 through their TADs

The thermomorphogenetic defect of *med14* and the requirement of MED14 for the thermal induction of many PIF4 direct targets suggest that it may be the Mediator tail subunit that communicates with PIF4 and/or HMR during the transcriptional activation of those thermoresponsive genes. To test this hypothesis, we investigated the interactions between MED14 and PIF4/HMR. We first expressed PIF4 and HMR as Glutathione S-transferase (GST) fusion proteins in *E. coli* and used them as baits to test their interactions with in-vitro transcribed/translated HA-MED14. Both GST-HMR and GST-PIF4 appear to interact strongly with HA-MED14 (Figure 4A).

**Figure 4.**
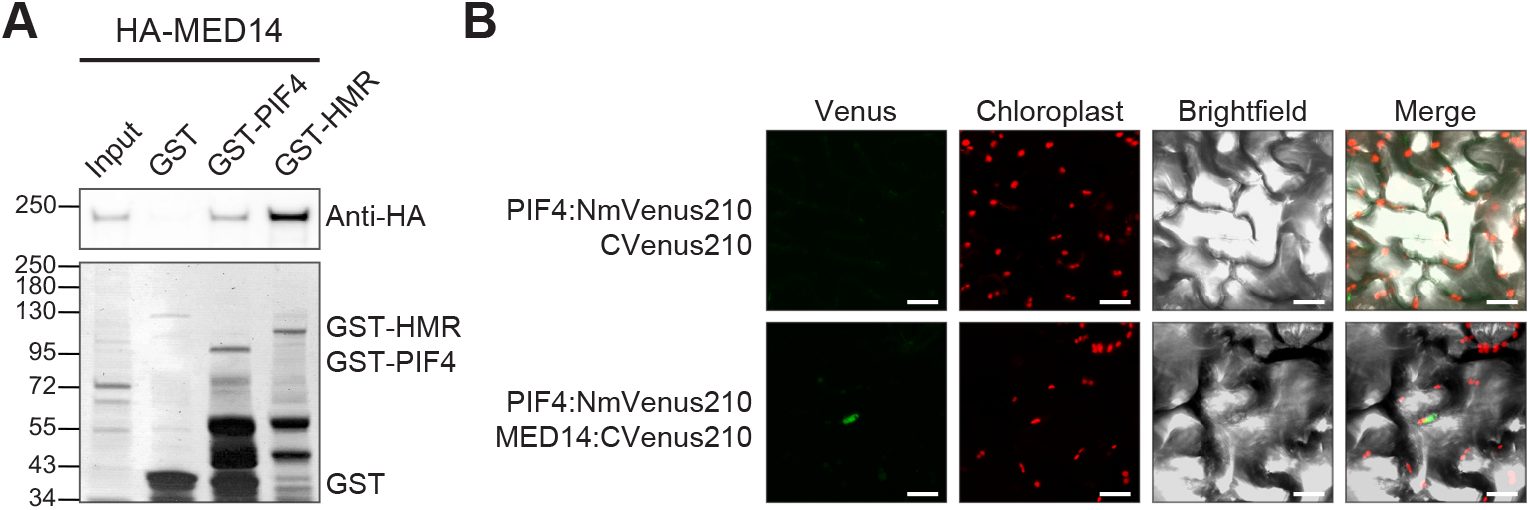
MED14 physically interacts with PIF4 and HMR. (A) GST pull-down assays. GST-tagged full-length PIF4 and HMR were used as baits to pull down in vitro translated HA-tagged MED14. The upper panel is an immunoblot using anti-HA antibodies showing the bound and input fractions of HA-MED14; the lower panel is a Coomassie Blue-stained SDS-PAGE gel showing immobilized GST and GST-tagged PIF4 and HMR. (B) Bimolecular fluorescence complementation assays. PIF4 fused with NmVenus210 (PIF4:NmVenus210) and MED14 fused with cVenus210 (MED14:CVenus210) were co-expressed in *Nicotiana benthamiana* through agroinfiltration. PIF4:NmVenus210 and CVenus210 were co-expressed as a control. The optical density of each *Agrobacterium* strain was 0.03 and images were captured 48 h after infiltration. The fluorescence signal (Venus) in the nucleus indicates the interactions between the two proteins tested in each agroinfiltration. The Chloroplast channel is to show that the Venus signal was not caused by the chlorophyll autofluorescence. Scale bars, 20 μm.

We further confirmed the interactions between PIF4 and MED14 using Bimolecular fluorescence complementation (BiFC) assays. When expressed in *Nicotiana benthamiana*, PIF4-NmVen210 interacted with MED14-cVen210 but not with cVen210 alone (Figure 4B). We did not observe the yellow fluorescent signal when co-expressing HMR-NmVen210 and MED14-cVen210, which was probably due to the fact that HMR is a nuclear/plastidial dual-localized protein, and its strong plastid transit peptide limits the abundance of HMR proteins in the nucleus (Chen et al., 2010; Nevarez et al., 2017). Taken together, the results from the GST pull-down assays and the BiFC assays indicate that MED14 may be physically associated with PIF4 and HMR.

### The association of MED14 with thermo-induced genes requires PIF4 and HMR

To elucidate the molecular mechanisms by which PIF4 and HMR recruit MED14 to the thermoresponsive PIF4 targets for transcriptional activation, we compared the association of MED14 with representative PIF4 targets in the presence and absence of functional PIF4 or HMR using ChIP-qPCR assays. The *35S::MED14-HA/med14* transgenic lines were crossed with *pif4-2* and *hmr-22*, respectively. The homozygous F3 progenies of *MED14-HA/pif4-2* and *MED14-HA/hmr-22* were used to compare MED14 occupancy across *IAA29* and *IAA19*, two PIF4/HMR/MED14-dependent thermoresponsive PIF4 direct targets, with that in *MED14-HA* after a 6-hour warm temperature treatment. The ChIP-qPCR results showed that MED14-HA is highly associated with the G-box (CACGTG)-containing regions in both *IAA29* and *IAA19*, confirming the physical presence of MED14 in the promoters of these two PIF4 targets (Supplemental Figure S4). Interestingly, this association was significantly reduced at the G-box-containing regions in the *pif4-2* or *hmr-22* mutant background (Figure 5A-D). We further crossed *MED14-HA/hmr-22* with *rcb-101*, a suppressor of *hmr-22* that can rescue *hmr-22*’s defects in thermomorphogenetic PIF4 activity and protein stability (Qiu et al., 2021). The association of MED14 with the G-box-containing regions in both *IAA29* and *IAA19* were rescued in the *MED14-HA/hmr-22*/*rcb-101* seedlings (Figure 5C, D). These data indicate that MED14 may be recruited by PIF4 and HMR to the promoters of growth-related PIF4 targets for their thermo-induced expression.

**Figure 5.**
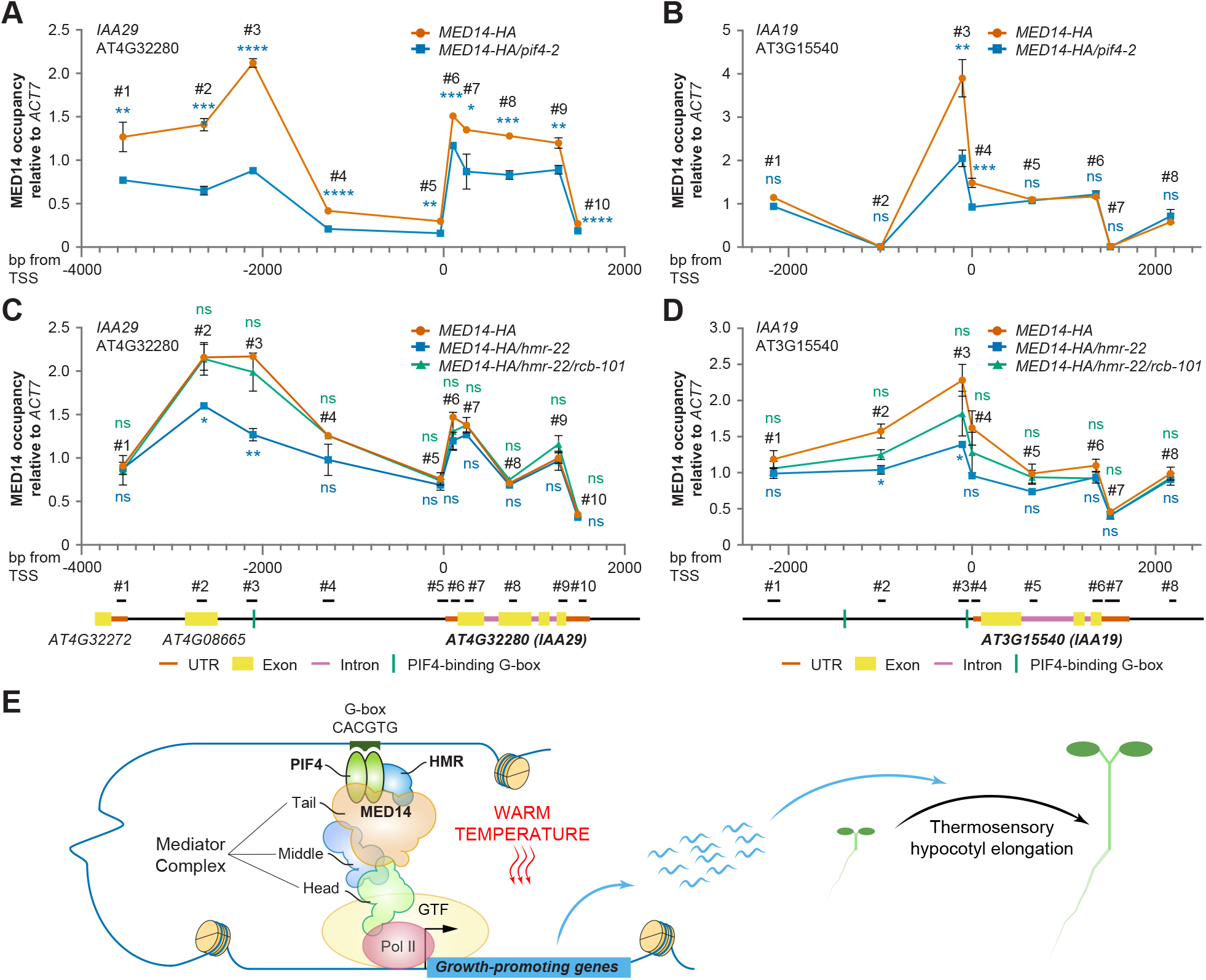
The association of MED14 with thermo-induced PIF4 direct targets is dependent on PIF4 and HMR. ChIP-qPCR experiments compare MED14 occupancy across *IAA29* (A and C) or *IAA19* (B and D) in the presence and absence of functional PIF4 (A and B) or HMR (C and D). Four-day-old 20 °C-grown *MED14-HA*, *MED14-HA/pif4-2*, *MED14-HA/hmr-22*, and *MED14-HA/hmr-22/rcb-101* were treated at 27 °C for 6 hours. HA antibody was used to assay MED14-HA association with each gene. Values are mean ± SD from two independent samples, with data presented as the ratio of MED14-HA at *IAA*/input to MED14-HA at *ACT7* (±1912)/input. Student t-tests were used to compare the values between MED14-HA and other lines. ****, *p* < 0.0001; ***, *p* < 0.001; **, *p* < 0.01; *, *p* < 0.05; ns, not significant (*p* ≥ 0.05). The exact position of each PCR fragment is shown in the illustrations at the bottom and the primers are listed in Supplemental Table S4. The transcription start site (TSS) is the first nucleotide of the 5’UTR, as defined by the Araport11 genome annotation, released June 2016. (E) Schematic model for MED14 recruitment by PIF4 and HMR during the thermo-induced transcription of growth-promoting PIF4 target genes, the products of which subsequently promote thermosensory hypocotyl elongation. Pol II, RNA polymerase II; GTF, general transcription factors. The black arrow indicates the TSS. The size and proportions of the depicted components do not reflect their actual dimensions.

## Discussion

Transcriptional regulation of thermoresponsive genes is one of the earliest and principal steps in plant thermomorphogenesis. We previously discovered that PIF4 and its coactivator HMR promote thermomorphogenetic hypocotyl growth in the daytime by activating the transcription of key growth-promoting genes (Qiu et al., 2019; Qiu et al., 2021). In this study, we further demonstrate that PIF4/HMR-mediated transcriptional regulation is achieved by recruiting the Mediator complex through their direct interactions with MED14 (Figure 5E).

The Mediator complex is required for virtually all Pol II-mediated transcription in eukaryotes (Jeronimo and Robert, 2017; Soutourina, 2018). As a crucial component of the Mediator complex, MED14 has long been known to play important roles in the transcription of key genes involved in various developmental processes and stress responses, including cell proliferation, shoot apical meristem development, trichome papillae development, salicylic acid-, methyl jasmonate-, and ethylene-mediated plant immune responses, abscisic acid-dependent drought responses, as well as cold and heat stress responses (Autran et al., 2002; Zhang et al., 2013; Hemsley et al., 2014; Wang et al., 2016; Fornero et al., 2019; Ohama et al., 2020; Lee et al., 2021a). In particular, MED14 works with MED16 and MED2 in modulating cold-responsive genes regulated by the AP2 transcription factors C-repeat binding factors (CBFs), and operates with MED17 in activating heat stress-inducible genes by assisting HsfA1, a heat shock transcription factor (HSF) that functions as a master regulator of the heat stress response (Hemsley et al., 2014; Ohama et al., 2020). We show here that MED14’s function is not limited to plant responses to extreme cold or hot temperatures, rather, it is also indispensable for the transcription of growth-related genes involved in moderately elevated temperature responses (Figure 1-3).

In higher eukaryotes, Mediator contains at least 7 tail components, namely MED2/29/32, MED3/27, MED5/24/33, MED15, MED16, MED23, and MED25 (Buendía-Monreal and Gillmor, 2016; Dolan and Chapple, 2017; Soutourina, 2018). Although originally assigned to the tail module (Dotson et al., 2000), MED14 was later found to play a more complex and critical role in both basal and activated transcription. For example, the cryo-electron microscopy (Cryo-EM) structure of *Schizosaccharomyces pombe* Mediator revealed that yeast MED14 functions as a backbone that connects the head, middle, and tail modules (Tsai et al., 2017). Likewise, *Mus musculus* MED14 is also centrally positioned, which enables inter-module interactions and makes it heavily involved in tail interactions (Zhao et al., 2021). Although it has not been definitively determined, Arabidopsis MED14 may possess similar structural features and play equally important roles in bridging all three modules as well as interacting with specific transcription factors, as suggested here by our data showing that MED14 physically interacts with activators PIF4 and HMR for thermoresponsive gene expression (Figure 4).

Nevertheless, the precise molecular details underlying the PIF4/HMR-MED14 interaction and the subsequent transcriptional activation of thermo-induced genes have yet to be revealed. Recent studies revealed that a variety of mammalian and yeast transcription factors form phase-separated condensates with Mediator through diverse TADs and that the formation of such condensates is associated with gene activation (Boija et al., 2018; Sabari et al., 2018). The phase-separating capacity of TADs and Mediator subunits is endowed by intrinsically disordered regions (IDRs), low-complexity protein segments that lack a defined secondary structure. In most eukaryotes, more than 80% of transcription factors and 75% of Mediator subunits possess extended (≥30 residues) IDRs (Liu et al., 2006; Tóth-Petróczy et al., 2008). Similarly, numerous plant transcription factors and Mediator subunits are predicted to contain at least one IDR (Nagulapalli et al., 2016; Salladini et al., 2020). In particular, the bHLH transcription factor family, to which PIF4 belongs, is hypothesized to have a general ability to undergo spontaneous liquid-liquid phase separation in transcription regulation by their IDRs (Tarczewska and Greb-Markiewicz, 2019; Salladini et al., 2020). Both PIF4 and HMR possess TADs that are potentially intrinsically disordered and thus could contribute to interactions with the Mediator complex. PIF4’s TAD contains a variant of the ΦxxΦΦ motif (Φ indicates a bulky hydrophobic residue and x is any other amino acid) flanked by multiple acidic residues, a structural feature that resembles the activator motifs in other activators such as mammalian p53 and yeast Gcn4 (Yoo et al., 2021). Mutating the ΦxxΦΦ motif reduced PIF4 transcriptional activity in yeast (Yoo et al., 2021). HMR possesses an acidic type 9aaTAD, the mutation of which drastically reduces its transactivation activity (Qiu et al., 2015) and affects MED14 recruitment to the promoter of PIF4-regulated thermo-induced genes (Figure 5). Future research will focus on unraveling the interaction mechanism between PIF4/HMR and MED14.

## Methods

### Plant materials and growth conditions

All the Arabidopsis mutants used in this study are in the Col-0 background. Mediator mutants were obtained from the Arabidopsis Biological Resource Center and the detailed list is shown in Supplemental Table S1. Homozygosity was confirmed by PCR before the seeds were used for phenotyping. Other Arabidopsis mutants, including *pif4-2* (SAIL_1288_E07) and *hmr-22*, were previously described (Leivar et al., 2008; Qiu et al., 2015; Qiu et al., 2019). Seeds were briefly rinsed with 70% ethanol and surface sterilized with bleach (3% sodium hypochlorite) for 10 min before being plated on half-strength Murashige and Skoog (1/2 MS) media supplemented with Gamborg’s vitamins (MSP0506, Caisson Laboratories, North Logan, UT), 0.5 mM MES (pH 5.7), and 0.8% (w/v) agar (A038, Caisson Laboratories, North Logan, UT). Seeds were stratified in the dark at 4 °C for 3-5 days to synchronize germination before treatment under specific light and temperature conditions in LED chambers (Percival Scientific, Perry, IA). Fluence rates of light were measured using an Apogee PS200 spectroradiometer (Apogee Instruments Inc., Logan, UT).

### Hypocotyl measurements

Seedlings were grown at either 20 °C or 27 °C under continuous R light (50 μmol m^−2^ s^−1^) for 96 hours. More than thirty seedlings from each line were placed on transparency film paper and scanned using an Epson Perfection V700 photo scanner. Hypocotyl length was measured using NIH ImageJ software (http://rsb.info.nih.gov/nih-image/). The percent increase in the hypocotyl length of each line was calculated as the percentage of the increase in hypocotyl length at 27 °C compared with that at 20 °C. The relative response of a mutant is defined as the percentage of its PI value or temperature response relative to that of Col-0. At least three replicates were used to calculate the mean and standard deviation of each relative response. Violin and box plots were generated using Prism 9 (GraphPad Software, San Diego, CA).

### Plasmid constructions

All PCR reactions were performed using the Q5 High-Fidelity DNA Polymerase and the ligation reactions with the NEBuilder HiFi DNA Assembly Master Mix (New England Biolabs Inc., Ipswich, MA). To generate the binary vector for making *MED14* transgenic lines, the full-length coding sequence (CDS) of *MED14* was fused with the sequence of a *(PT)_4_P linker* and 3 copies of an *HA-tag* when being cloned into the binary vector pCHF1 between SacI and BamHI. The bait vectors used in GST pull-down assays were either made previously (Qiu et al., 2015) or generated by cloning the full-length CDS of *PIF4* into pET42b vectors between BamHI and HindIII, and prey vectors were constructed by cloning the full-length CDS of *MED14* into the pCMX-PL2-NterHA vector between EcoRI and BamHI. Vectors used for the Bimolecular fluorescence complementation assay were generated by cloning the full-length CDS of *PIF4* and *HMR* into MCS I of pDOE-01 (ABRC stock# CD3-1901) between NcoI and SpeI, followed by cloning the full-length CDS of *MED14* into MCS III between KfII and AatII. All the primers used for plasmid constructions are listed in Supplemental Table S2.

### Generation of transgenic lines

To generate the *35S::MED14-HA/med14* transgenic lines, the *med14* mutant was transformed with the above-described *pCHF1-MED14-(PT)_4_P-3×HA* plasmid using the *Agrobacterium*-mediated floral dip method. The transformants were selected on the 1/2 MS medium containing 100 μg/ml gentamycin. At least 10 independent lines that segregated approximately 3:1 for gentamycin-resistance in the T2 generation were identified and two of these lines were selected for this study on the basis of the transgene expression level. For all experiments, T3 self-progenies of homozygous T2 plants were used.

### RNA extraction and quantitative reverse transcription PCR (RT-qPCR)

Total RNA was extracted from 50-100 mg seedlings using the Quick-RNA Miniprep Kit with on-column DNase I digestion (Zymo Research, Irvine, CA). cDNA was synthesized with 2–2.5 μg total RNA using the Invitrogen SuperScript III Reverse Transcriptase and the Oligo(dT)_20_ primer (Thermo Fisher Scientific, Waltham, MA). For RT-qPCR, cDNA diluted in nuclease-free water was mixed with FastStart Universal SYBR Green Master (MilliporeSigma, Burlington, MA) and gene-specific primers (Supplemental Table S3). RT-qPCR reactions were performed in triplicate with a Qiagen Rotor-Gene Q 5Plex real-time cycler (Qiagen, Germantown, MD). Transcript levels of each gene were calculated relative to that of *PP2A*. A standard curve was performed for each gene to determine the linear range, efficiency, and reproducibility of the qPCR assay. Bar charts were generated using Prism 9 (GraphPad Software, San Diego, CA).

### RNA-seq and data analysis

RNA was isolated as described above. The RNA concentration and purity were determined spectrophotometrically using the Nanodrop 2000 instrument (Thermo Fisher Scientific, Waltham, MA), and assessing the continuous spectrum (190 nm to 840 nm) generated by the instrument for each sample. The quality of the RNA samples was evaluated on the Agilent 2100 Bioanalyzer (Agilent Technologies Inc., Santa Clara, CA).

Sequencing libraries were generated with 300 ng to 1 μg of RNA using the TruSeq Stranded mRNA Sample Preparation Kit (Illumina, San Diego, CA). The libraries were assessed for size and purity using the Agilent 2100 Bioanalyzer and quantitated by qPCR using a KAPA Biosystems Library Quantitation kit (Roche, Pleasanton, CA). Libraries were normalized, pooled, and diluted to a loading concentration of 1.8 pM, loaded on an Illumina High-Output Flow Cell, and sequenced on the Illumina NextSeq 500 instrument. The sequencing run generated ~16 million 150 bp paired-end reads per sample, and all run metrics, including Q-Score distributions, cluster densities, and total sequence yields, were within the recommended parameters.

Data analysis was performed using the Qiagen CLC Genomics Workbench (CLCGWB) version 21 software. The reads were mapped to the A. thaliana reference genome (TAIR10/Araport11 genome release), using default parameters with a strand-specific alignment protocol. The mapping report indicated that all metrics fell within the recommended parameters, with >90% of the reads per sample mapping to the reference genome. The mapping results generated values for Total Counts, Transcripts per Million reads, as well as Fragments Per Kilobase of exon per Million reads mapped (FPKM) for each gene. The “Differential Expression for RNA-Seq” tool in CLCGWB software was used, with default parameters, to identify differentially expressed genes across samples. Genes that had a fold-change of ≥2, an FDR-corrected p-value of <0.001, and a Maximum Group Mean FPKM of >1 (i.e., the maximum of the average FPKM across the two groups in each statistical comparison) were considered to be significantly differentially expressed. The list of differentially expressed genes obtained in each statistical comparison is shown in Supplemental Dataset 1. Venn diagrams were generated using the Bioinformatics and Evolutionary Genomics website (https://bioinformatics.psb.ugent.be/webtools/Venn/). Hierarchical cluster analysis was performed with the Morpheus software tool available at the Broad Institute website (https://software.broadinstitute.org/morpheus/). Gene Ontology (GO) enrichment analyses were performed using the PANTHER classification tool available at the TAIR website (https://www.arabidopsis.org/tools/go_term_enrichment.jsp) or the GeneCodis 4 program (https://genecodis.genyo.es/) (García-Moreno et al., 2021). For comparison of data with published datasets, the appropriate FastQ files available in the NCBI-GEO database were imported into CLCGWB software, and all analyses were conducted in a similar manner to those described above. For the published datasets that utilized a 3’TagSeq protocol, mapping was performed using the 3’-Sequencing option in the CLCGWB alignment parameters.

RNA-Seq data accession number. The RNA-Seq analysis data described in this article are accessible through accession no. GSE196969 at NCBI’s Gene Expression Omnibus database.

### Protein extraction and immunoblots

For immunoblots, total protein was extracted from 4-day-old seedlings as previously described (Qiu et al., 2019; Qiu et al., 2021). Briefly, 200 mg seedlings were freshly homogenized using a BeadBug microtube homogenizer (Benchmark Scientific Inc., Sayreville, NJ) in three volumes (mg/μL) of extraction buffer containing 100 mM Tris-HCl pH 7.5, 100 mM NaCl, 5 mM EDTA, 5% SDS, 20% glycerol, 20 mM DTT, 40 mM β-mercaptoethanol, 2 mM phenylmethylsulfonyl fluoride, 40 μM MG115 (Apexbio Technology LLC, Houston, TX), 40 μM MG132 (Cayman Chemical, Ann Arbor, MI), 40 μM bortezomib (MilliporeSigma, Burlington, MA), 10 mM N-ethylmaleimide (Thermo Fisher Scientific, Waltham, MA), 1× phosphatase inhibitor cocktail 2 (Thermo Fisher Scientific, Waltham, MA), 1× phosphatase inhibitor cocktail 3 (MilliporeSigma, Burlington, MA), 1× EDTA-free protease inhibitor cocktail (MilliporeSigma, Burlington, MA), and 0.01% bromophenol blue. Samples were boiled for 10 min and centrifuged at 16,000 × g for 10 min. The supernatant was immediately used for immunoblots or stored at −80 °C until use.

For immunoblots, cleared protein samples were separated via SDS-PAGE, transferred to nitrocellulose membranes, probed with the indicated primary antibodies, and then incubated with 1:5000 dilution of horseradish peroxidase-conjugated goat anti-rabbit or anti-mouse secondary antibodies (Bio-Rad Laboratories, 1706515 for anti-rabbit and 1706516 for anti-mouse). Primary antibodies, including monoclonal mouse anti-HA antibodies (MilliporeSigma, H3663), polyclonal rabbit anti-HMR antibodies (Chen et al., 2010), polyclonal goat anti-PIF4 antibodies (Agrisera, AS16 3955), and polyclonal rabbit anti-RPN6 antibodies (Enzo Life Sciences, BMLPW8370-0100) were used at 1:1000 dilution. Signals were detected via chemiluminescence using a SuperSignal kit (Thermo Fisher Scientific, Waltham, MA) and an Azure C600 Advanced Imaging System (Azure Biosystems, Dublin, CA).

### Chromatin immunoprecipitation

ChIP assays were performed as previously described with modifications (Wu et al., 2016). Seedlings grown on ½ MS medium in continuous red light (50 μmol m^−2^ s^−1^) at 20 °C for 96 h were treated at 27 °C for 6 h before collecting. About 200 mg seedlings were cross-linked with 1% formaldehyde in 1× PBS for 2× 15 min by vacuum infiltration, followed by the addition of glycine to 125 mM with another 10 min of vacuum infiltration. Fixed seedlings were dried on a stack of paper towels after being washed with 4× 10 ml cold PBS. Samples were ground into a fine powder in liquid nitrogen and resuspended in Nuclear Isolation Buffer (20 mM Hepes, pH 7.6, 0.25 M sucrose, 5 mM KCl, 10 mM MgCl_2_, 40% glycerol, 0.25% Triton X-100, 0.1 mM PMSF, 0.1% β-mercaptoethanol, 1× complete EDTA-free protease inhibitor cocktail (MilliporeSigma, Burlington, MA). Nuclei were enriched by filtering the lysate through two layers of Miracloth and centrifugation at 3,000 × g for 10 min at 4 °C. Nuclear pellets were resuspended in Nuclei Lysis Buffer (20 mM Tris-HCl, pH 7.5, 100 mM NaCl, 2.5 mM EDTA, 10% glycerol, 1% Triton X-100, 1× protease inhibitor cocktail) and sonicated with 25 cycles of 30-s-long pulses (30-s intervals) using a Bioruptor Plus (Diagenode, Denville, NJ). Nuclear lysates were cleared with 2 rounds of centrifugation at 13,000 × g for 5 min at 4 °C. Immunoprecipitation was performed by incubating 30 μL of Dynabeads Protein G (Thermo Fisher Scientific, Waltham, MA), 2.5 μg of anti-HA tag antibody (Abcam, Waltham, MA), and 1 mL of diluted chromatin (containing 100 μL of sonicated chromatin) at 4 °C for 4 h. Beads were washed for 2 × 15 min with low-salt wash buffer (20 mM Tris-HCl, pH 7.5, 150 mM NaCl, 2.5 mM EDTA, 0.05% SDS, 1% Triton X-100), for 2 × 15 min with high-salt wash buffer (20 mM Tris-HCl, pH 7.5, 500 mM NaCl, 2.5 mM EDTA, 0.05% SDS, 1% Triton X-100), for 1 × 15 min with LiCl Wash Buffer (20 mM Tris-HCl, pH 7.5, 250 mM LiCl, 1 mM EDTA, 0.5% Na-deoxycholate, 0.5% Nonidet P-40), and for 1 × 15 min with TE buffer (10 mM Tris-HCl, pH 7.5, 1 mM EDTA). 10% (wt/vol) chelex resin (Bio-Rad, Hercules, CA) was mixed with the beads and immunoprecipitated DNA was eluted after reverse cross-linking by boiling at 95 °C for 10 min, followed by treatment with 40 μg of proteinase K for 1 h at 55 °C. Eluted DNA was recovered using the ChIP DNA Clean & Concentrator kit (Zymo Research, Irvine, CA) and used for qPCR reactions. Primers for ChIP-qPCR are listed in Supplemental Table S4. Values are means ± SD from two independent samples; data are presented as a ratio of (MED14-HA *IAA*/input *IAA*) to (MED14-HA at internal control/input at internal control) to correct for tube-to-tube variation (the MED14-HA level at position +1912 in *Actin7* was used as an internal control for normalization).

### GST pull-down

GST pull-down assays were performed as described previously (Qiu et al., 2015; Qiu et al., 2021). The pET42b/PIF4 and pET42b/HMR vectors were constructed as described above and expressed as GST-PIF4 and GST-HMR fusion proteins in the *E. coli* strain BL21-CodonPlus (DE3) (Agilent Technologies). The pCMX-PL2-NterHA/MED14 vector was constructed as described above and expressed as HA-MED14 proteins using the TNT T7 Coupled Reticulocyte Lysate System (Promega, Madison, WI). HA-MED14 prey proteins were incubated with the affinity-purified GST-PIF4 or GST-HMR bait proteins immobilized on glutathione Sepharose 4B beads (GE Healthcare, Chicago, IL) at 4 °C for 2 h. Beads were washed four times with E buffer (50 mM Tris-HCl, pH 7.5, 100 mM NaCl, 1 mM EDTA, 1 mM EGTA, 1% DMSO, 2 mM DTT, 0.1% Nonidet P-40). Bound proteins were eluted by boiling in 2× SDS loading buffer and used in subsequent SDS-PAGE and immunoblotting. Input and immunoprecipitated HA-MED14 prey proteins were detected using goat anti-HA polyclonal antibodies (GenScript, Piscataway, NJ). The amounts of GST-PIF4 and GST-HMR bait proteins were visualized by staining the SDS-PAGE with Coomassie Brilliant Blue.

### Agroinfiltration-based Bimolecular fluorescence complementation (BiFC)

The pDOE-01/PIF4 and pDOE-01/PIF4/MED14 vectors were constructed as shown above. The BiFC assays were performed as described (Gookin and Assmann, 2014). The constructed pDOE-01 vectors were transformed into GV3101 *Agrobacterium* electrocompetent cells and selected on 25 μg/ml rifampicin, 20 μg/ml gentamycin, and 50 μg/ml kanamycin at 29 °C for 2-3 days. Several colonies of positive transformants were harvested from the plate, inoculated to LB media containing the above-mentioned antibiotics, and cultured at 29 °C for 2 h with 100 μM acetosyringone (MilliporeSigma, Burlington, MA). The *Agrobacterium* cells were washed with and diluted to an OD_600_ of 0.03 in the infiltration buffer (10 mM MES-KOH, pH 5.6, 10 mM MgCl_2_, and 100 μM acetosyringone).

*Nicotiana benthamiana* plants were grown in General Purpose Pro-Mix BX (Premier Tech Horticulture, Quakertown, PA) in a growth room with 150 μmol/m^2^/sec white light, 14-h light/10-h dark cycles, and a temperature range between 21-24 °C. BiFC control and test infiltrations were performed on three plants each. Fluorescence signals were examined 48-72 hours after agroinfiltration using a Leica SP8 Inverted Confocal Microscope (Leica Microsystems Inc., Buffalo Grove, IL). Excitation and emission wavelengths for mVenus were 488 nm and 500-550 nm, respectively.

## Accession Numbers

Accession numbers are as described by TAIR (https://www.arabidopsis.org) as follows: PHYTOCHROME-INTERACTING FACTOR 4 (PIF4), AT2G43010; HEMERA (HMR), AT2G34640; MED2/29/32, AT1G11760; MED3/27, AT3G09180; MED5A/33A, AT3G23590; MED5B/33B, AT2G48110; MED6, AT3G21350; MED7A, AT5G03220; MED7B, AT5G03500; MED8, AT2G03070; MED9, AT1G55080; MED10A, AT5G41910; MED10B, AT1G26665; MED14, AT3G04740; MED16, AT4G04920; MED17, AT5G20170; MED18, AT2G22370; MED19A, AT5G12230; MED22A, AT1G16430; MED22B, AT1G07950; MED23, AT1G23230; MED25, AT1G25540; MED28, AT3G52860; MED30, AT5G63480; MED31, AT5G19910; CYCLINC1-2 (CYCCA), AT5G48630; CYCLINC1-1 (CYCCB), AT5G48640; SERINE/THREONINE PROTEIN PHOSPHATASE 2A (PP2A), AT1G69960; YUCCA8 (YUC8), AT4G28720; INDOLE-3-ACETIC ACID INDUCIBLE 19 (IAA19), AT3G15540; INDOLE-3-ACETIC ACID INDUCIBLE 29 (IAA29), AT4G32280; SULFOTRANSFERASE 2A (ST2A), AT5G07010; ARABIDOPSIS PP2C CLADE D 7 (APD7), AT5G02760; AUXIN-REGULATED GENE INVOLVED IN ORGAN SIZE (ARGOS), AT3G59900; RING, AT1G24580; SERINE CARBOXYPEPTIDASE-LIKE 31 (SCPL31), AT1G11080; METHYL ESTERASE 18 (MES18), AT5G58310; ACTIN 7 (ACT7), AT5G09810.

## Supplemental Data

Supplemental Figure S1. Thermomorphogenetic hypocotyl responses in mediator mutants.

Supplemental Figure S2. Comparison of thermoresponsive transcriptomes in wild-type seedlings from different studies.

Supplemental Figure S3. Analysis of PIF4/HMR/MED14-dependent thermoresponsive genes.

Supplemental Figure S4. MED14 is associated with thermo-induced PIF4 direct targets at warm temperatures.

Supplemental Table S1. List of T-DNA insertion lines and genotyping primers used in this study.

Supplemental Table S2. Primers used for plasmid construction.

Supplemental Table S3. RT-qPCR primers for the genes examined in this study.

Supplemental Table S4. ChIP-qPCR primers for the genes examined in this study.

Supplemental Dataset 1: Lists of SSTF genes regulated by warm temperatures in Col-0, *pif4-2*, *hmr-22*, and *med14*.

Supplemental Dataset 2: Lists of thermo-responsive genes in two other reports.

Supplemental Dataset 3: Lists of PIF4/HMR/MED14-dependent thermo-responsive genes.

Supplemental Dataset 4: Lists of potential PIF4/HMR/MED14-dependent, thermo-responsive PIF4 direct targets.

## Data Availability Statement

The original contributions presented in the study are included in the article/Supplementary Material, further inquiries can be directed to the corresponding author. The datasets presented in this study can be found in online repositories. The names of the repository/repositories and accession number(s) can be found at NCBI’s Gene Expression Omnibus database (https://www.ncbi.nlm.nih.gov/geo/), GSE196969.

## Acknowledgments

We would like to thank Dr. Gregg Roman and Dr. Ruofan Cao in the Imaging Research Core of GlyCORE at the University of Mississippi for their assistance with the usage of the Leica SP8 Inverted Confocal Microscope. We also thank all the lab members, especially Ranjeeta Odari and Anupa Wasti, for their critical comments and suggestions regarding the manuscript.

## Author Contributions

YQ and AB conceived the original research plan; YQ, SRB, and AKA supervised the experiments; AB, JX, KFG, EEB, CAG, CHC, AKA, and YQ performed the experiments; AB, JX, SRB, AKA, and YQ analyzed the data; YQ wrote the article with the contributions from all authors; all authors approved the submitted version.

